# Brain aging is faithfully modelled in organotypic brain slices and accelerated by prions

**DOI:** 10.1101/2022.02.06.479288

**Authors:** Yingjun Liu, Assunta Senatore, Silvia Sorce, Mario Nuvolone, Jingjing Guo, Zeynep H. Gümüş, Adriano Aguzzi

## Abstract

Mammalian models are essential for brain aging research. However, the long lifespan and limited amenability to genetic and pharmacological perturbations have hindered the use of mammals for dissecting aging-regulatory molecular networks and discovering new anti-aging interventions. To circumvent these limitations, we developed an ex vivo model system that faithfully mimics the aging process of the mammalian brain using cultured mouse brain slices. Genome-wide gene expression analyses showed that brain slices spontaneously upregulated senescence-associated genes over time and reproduced many of the transcriptional characteristics of aged brains. Treatment with rapamycin, a classical anti-aging compound, largely abolished the time-dependent transcriptional changes in brain slices. Using this model system, we discovered that prions drastically accelerated the development of age-related molecular signatures and the pace of brain aging. We confirmed this finding in mouse models and human victims of Creutzfeldt-Jakob disease. These data establish a novel, eminently tractable mammalian model of brain aging, and uncover a surprising acceleration of brain aging in prion diseases.

## Introduction

Advanced age is a strong risk factor for several chronic disorders affect multiple organs, including the brain (Niccoli and Partridge 2012). Therefore, treatments targeting the biological aging process may represent promising therapeutic interventions against many of these diseases. However, current mammalian models for aging research suffer from long lifespans and poor amenability for genetic and pharmacological perturbations (Brunet 2020), which limits their usefulness for dissecting aging-regulatory molecular networks and discovering new anti-aging interventions. Consequently, only a small number of molecular pathways relevant to human aging have been identified (Bitto,Wang et al. 2015). More versatile model systems are needed to advance our knowledge on aging of complex tissues in mammals including humans.

Previous studies have highlighted several hallmarks of biological aging, including inflammation, loss of proteostasis, dysregulated nutrient sensing, mitochondrial malfunction, and altered intercellular communication (Ferrucci, Gonzalez-Freire et al. 2020). Many of these aging hallmarks are also critical pathological events observed in brain disorders characterized by abnormal protein aggregation, such as prion diseases, a group of neurodegenerative disorders caused by misfolding and aggregation of the cellular prion protein (PrP^C^) into pathological isoforms called prions (Aguzzi and Liu 2017). Therefore, studying the aging process in protein-aggregation disorders of the brain may not only enhance our understanding of disease progression but also offer a unique opportunity to identify key drivers of biological brain aging.

All major pathological characteristics of prion diseases can be faithfully mimicked in both prion-inoculated animals and cultured organotypic cerebellar slices (COCS) (Falsig, Julius et al. 2008,Falsig, Sonati et al. 2012). Such cultures preserve the sophisticated cell-cell interactions present in vivo while gaining simplicity and experimental amenability, and represent a tantalizing biological system for investigating complex physiological and pathological processes of the brain. It is however unclear whether COCS experience biological aging akin to brains in vivo.

Here, we evaluated the suitability of COCS for modeling in vivo brain aging through genome-wide gene expression profiling, bioinformatics and machine learning, approaches widely used to measure and predict biological ages of various tissues (Jylhava, Pedersen et al. 2017, Fleischer, Schulte et al. 2018, Bulteau and Francesconi 2021, Meyer and Schumacher 2021). We found that COCS faithfully mimicked the in vivo molecular brain aging process, and demonstrated the physiological relevance of these findings through rapamycin treatment, establishing a novel, convenient, mammalian model system for brain aging research. Furthermore, we discovered that prions drastically accelerated the pace of brain aging in COCS, mouse models and human patients, providing a theoretical basis for exploiting “rejuvenating” therapies against prion diseases.

## Results

### Natural aging of COCS

We used RNA sequencing (RNAseq) to obtain genome-wide gene expression profiles of COCS maintained *ex vivo* for 12, 28, 42 or 56 days in the presence or absence of prions (**Fig. 1A**). To compare it with *in vivo*, similar RNAseq was performed for the cerebellum of adult C57BL/6J mice at 56 or 182 days post-intracerebral inoculation with normal brain homogenate (NBH) or brain homogenate containing prions (**Fig. 1A**). The global gene expression in the control COCS was highly similar to that in the cerebellum and hippocampus *in vivo*, but substantially different from the adult muscle (Borsch, Ham et al. 2021) (**Fig. 1B**), indicating that brain-specific gene expression was retained in COCS. The correlation of transcriptional patterns between the control COCS and adult brain tissues underwent a subtle monotonic decrease over time (**Fig. 1B**). Indeed, comparison of gene expression in the control COCS between 56 and 12 days identified 2736 differentially expressed genes (DEG) (**Supplementary fig. 1A**; **Supplementary table 1**).

**Figure 1.**
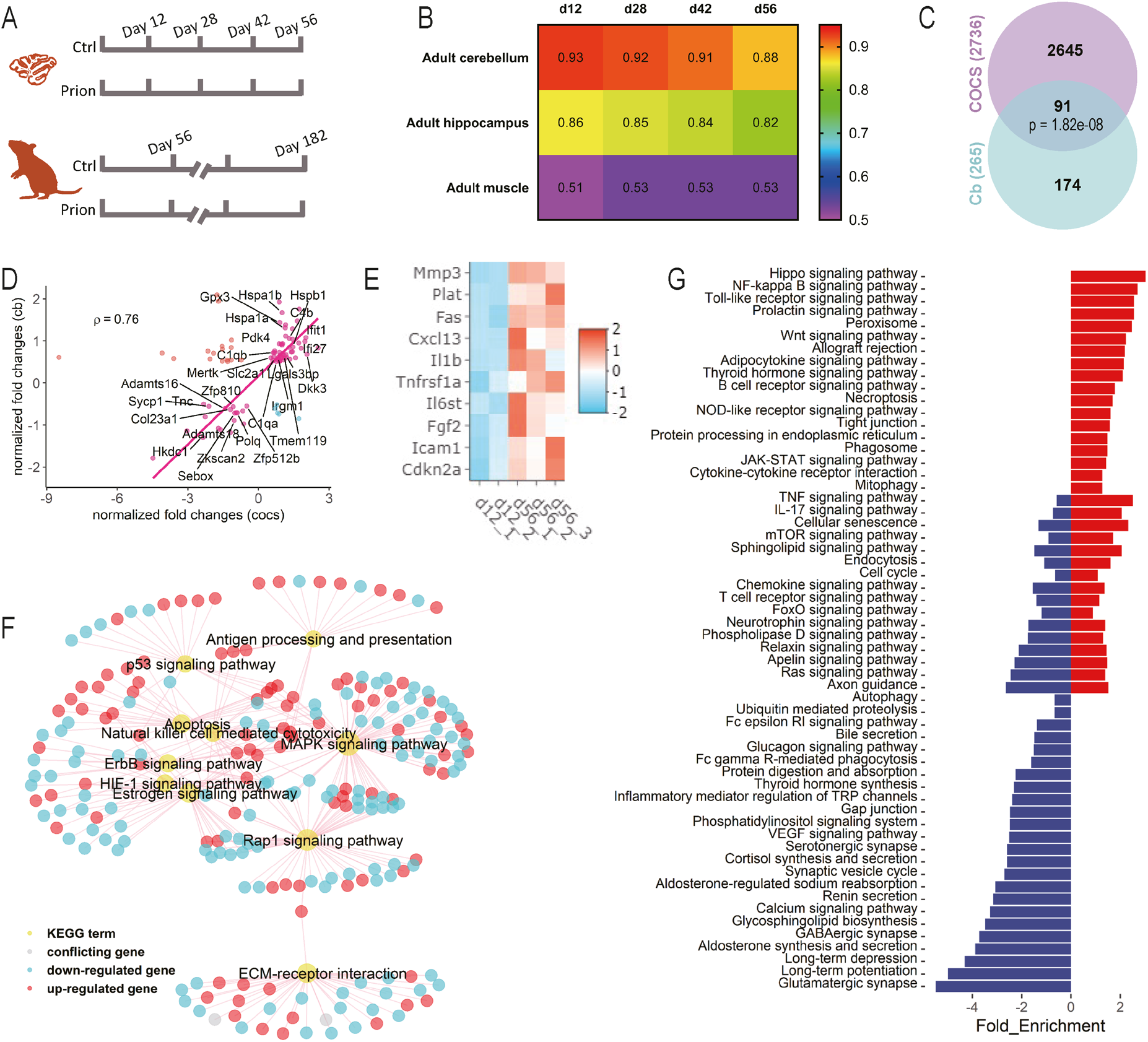
COCS recapitulate in vivo gene expression and undergo rapid aging. **A**. Design of the RNAseq experiments. **B**. Spearman’s correlation coefficient (ρ) of gene expression in COCS (12-56 days) with adult mouse cerebellum, hippocampus and muscle. X-axis indicates days ex vivo of COCS. **C**. Overlap between DEG of aged COCS and cerebellum (*p*: hypergeometric test). **D**. Changes of common DEG in aged COCS vs cerebella. Genes of interest are highlighted with their names. Genes show different directions of changes in the two conditions are highlighted in cyan or red. **E**. Heatmap showing the expression of cellular senescence genes, including p16 (Cdkn2a) and members of the senescence-associated secretory phenotype, in young (12 days) and old (56 days) COCS. **F**. Subset of common KEGG pathways that are significantly (FDR < 0.05) dysregulated in aged COCS and cerebellum. **G**. Additional KEGG pathways that are significantly (FDR < 0.05) enriched in the DEG of aged COCS. Negative and positive values of the x-axis indicate downregulation and upregulation during aging, respectively. Blue bars: downregulated pathways in aged COCS. Red bars: upregulated pathways in aged COCS.

To assess whether the time-dependent transcriptional changes in the control COCS mirror the physiology of brain aging in vivo, we examined whether similar alterations were detectable in the mouse cerebellum across ages. We identified 265 DEG in the cerebellar tissues of NBH-inoculated mice between 56 and 182 days (~4-9 months of age) (**Supplementary fig. 1B**; **Supplementary table 2**). Ninety-one out of the 265 DEG overlapped with those observed in the control COCS (**Fig. 1C**). Furthermore, 80% of the overlapping genes were altered in the same direction, and were highly correlated (ρ = 0.76, Spearman’s correlation) between the control COCS and in vivo (**Fig. 1D**). These genes were associated with microglia (e.g., *Tmem119*, *C1qa*, *C1qb* and *C4b*), inflammation (e.g., *Mertk*, *Irgm1*, *Ifit1*, *Ifi27* and *Lgals3bp*), protein stabilization (e.g., *Hspa1a*, *Hspa1b*, *Hspb1* and *Dkk3*),glucose metabolism (e.g., *Slc2a1*, *Pdk4*, *Gpx3* and *Hkdc1*), extracellular matrix (e.g., *Col23a1*, *Adamts16*, *Adamts18* and *Tnc*), cell proliferation and gene expression regulation (e.g., *Sycp1*, *Polq*, *Zkscan2*, *Sebox*, *Zfp512b* and *Zfp810*). In addition, a noteworthy overlap was identified between the DEG in the control COCS and more advanced brain-aging stages, such as the 24-month mouse hippocampus (Stilling, Benito et al. 2014) (**Supplementary fig. 2A-C**). Examination of the expression levels of cellular senescence genes, including the senescence marker p16 (*Cdkn2a*) and members of the senescence-associated secretory phenotype (e.g., *Mmp3* and *Icam1*), indicated a strong induction of cellular senescence in the control COCS at 56 days (**Fig. 1E**), recapitulating a hallmark of in vivo tissue aging (Yousefzadeh, Zhao et al. 2020).

To examine whether the transcriptional changes in the control COCS might be caused by the transient exposure to NBH, we analyzed ten randomly picked DEG (5 upregulated and 5 downregulated) and ten genes related to cellular senescence by quantitative RT-PCR. However, none of them showed any difference between the control and naïve (no NBH exposure) COCS at 12 and 56 days (**Supplementary fig. 3A-B)**, indicating that the time-dependent gene expression changes in the control COCS were mainly induced by the natural aging process.

Enrichment of DEG in aged COCS and in vivo for KEGG pathways identified 41 commonly altered biological processes (**Supplementary table 3**). These included antigen processing and presentation, p53 signaling, apoptosis, natural killer cell mediated cell toxicity, Mapk signaling, ErbB signaling, Hif1 signaling, estrogen signaling, Rap1 signaling and ECM-receptor interaction (**Fig. 1F**), all of which had been found to be overrepresented in aged brains (Frahm, Srivastava et al. 2017, Mattson and Arumugam 2018, Shavlakadze, Morris et al. 2019, Alique, Sanchez-Lopez et al. 2020, Paterson,Cumming et al. 2020). Additional aging-related pathways, such as NF-kB signaling, mTOR signaling, autophagy, cellular senescence and VEGF signaling, were found to be more profoundly altered in long-term cultured COCS than in 9-month cerebella (**Fig. 1G**), suggesting that the 56-day-old COCS represent a more advanced in vivo brain aging stage.

### Aging-modulatory effects of rapamycin in COCS

To further validate the physiological relevance of the time-dependent transcriptional changes in COCS to in vivo brain aging, we treated naïve COCS with rapamycin, which is often used in anti-aging paradigms, or with DMSO from day 12 to day 56, and examined the expression levels of cellular senescence genes by quantitative RT-PCR. Strikingly, we found that the transcriptional induction of all the cellular senescence-associated genes in aged COCS was drastically suppressed by rapamycin treatment (**Fig. 2A**). In addition, by applying elastic net-based machine learning (Zou and Hastie 2005) to our time-course RNAseq dataset of the control COCS, we identified 322 age-predicting genes (**Supplementary table 4**). Quantitative RT-PCR results demonstrated that rapamycin substantially inhibited the expression changes of all eight randomly picked upregulated age-predicting genes (**Fig. 2B**), and five out of eight randomly picked downregulated age-predicting genes (**Fig. 2B**), in the 56-day-old naïve COCS, indicating a global slowdown of the biological aging process upon rapamycin treatment. These data confirm that a biological process similar to that underlying in vivo brain aging drove the progressive transcriptional changes in COCS over time.

**Figure 2.**
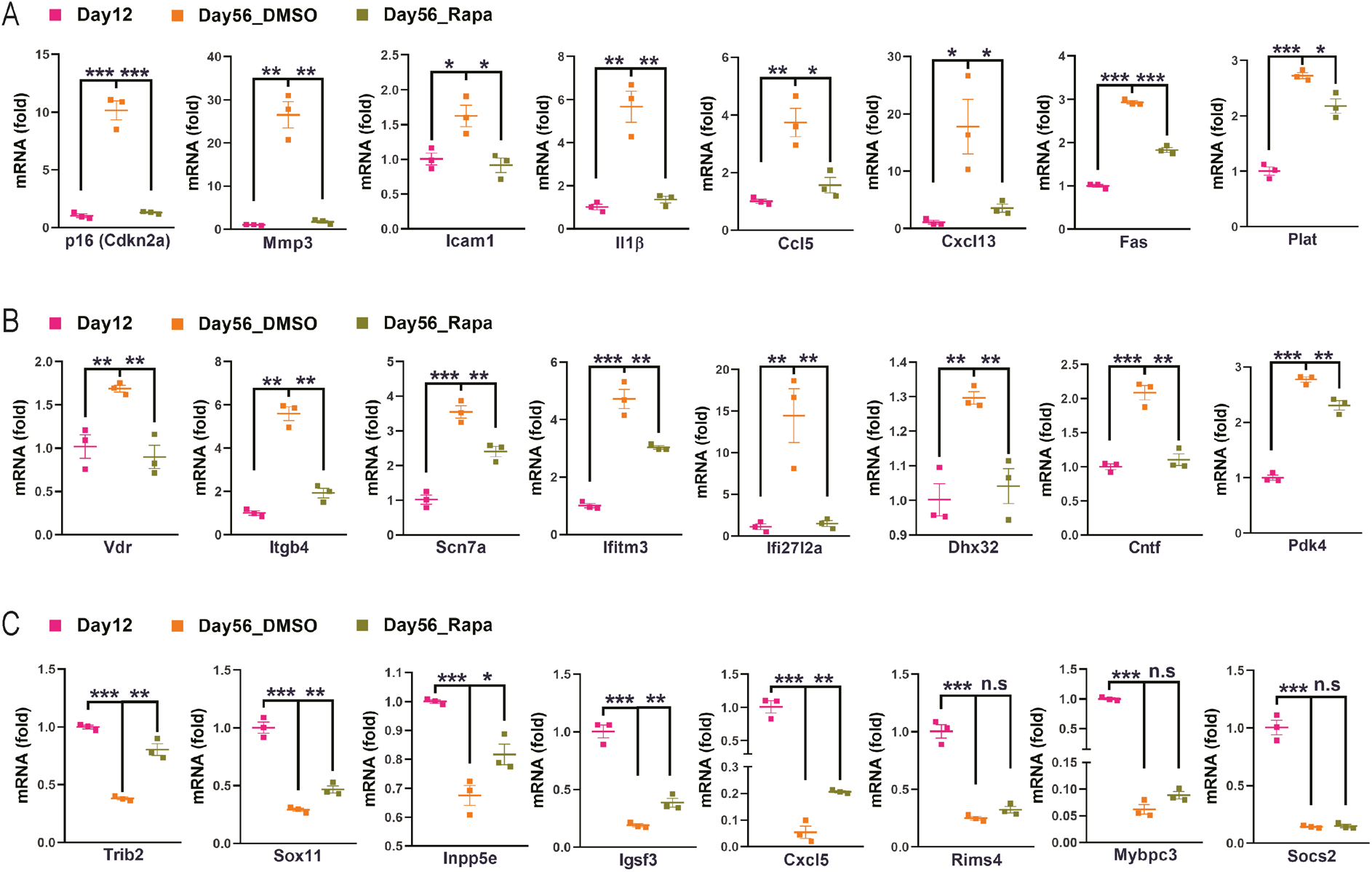
Aging-regulatory effects of rapamycin in COCS. **A**, Rapamycin (Rapa) decelerated the time-dependent upregulation of cellular senescence genes, including the senescence marker p16 (Cdkn2a) and members of the senescence-associated secretory phenotype. **B-C**, Rapamycin (Rapa) decelerated the time-dependent upregulation (B) and downregulation (C) of eight and five randomly picked age-predictive genes in COCS, respectively. *: p < 0.05, **: p < 0.01, ***: p < 0.001. n.s: not significant.

### Accelerated aging in prion-exposed COCS

To evaluate the power of our model system for identifying novel modifiers of brain aging, we next studied how the biological aging process develops in COCS exposed to prions. We compared the transcriptional changes in the prion-exposed COCS at 56 days with those induced by natural aging in the control COCS. Surprisingly, we found a large number of genes dysregulated in the aged COCS further changed their expression levels in the presence of prions (**Fig. 3A**). These results suggest that prions may alter the biological aging process.

**Figure 3.**
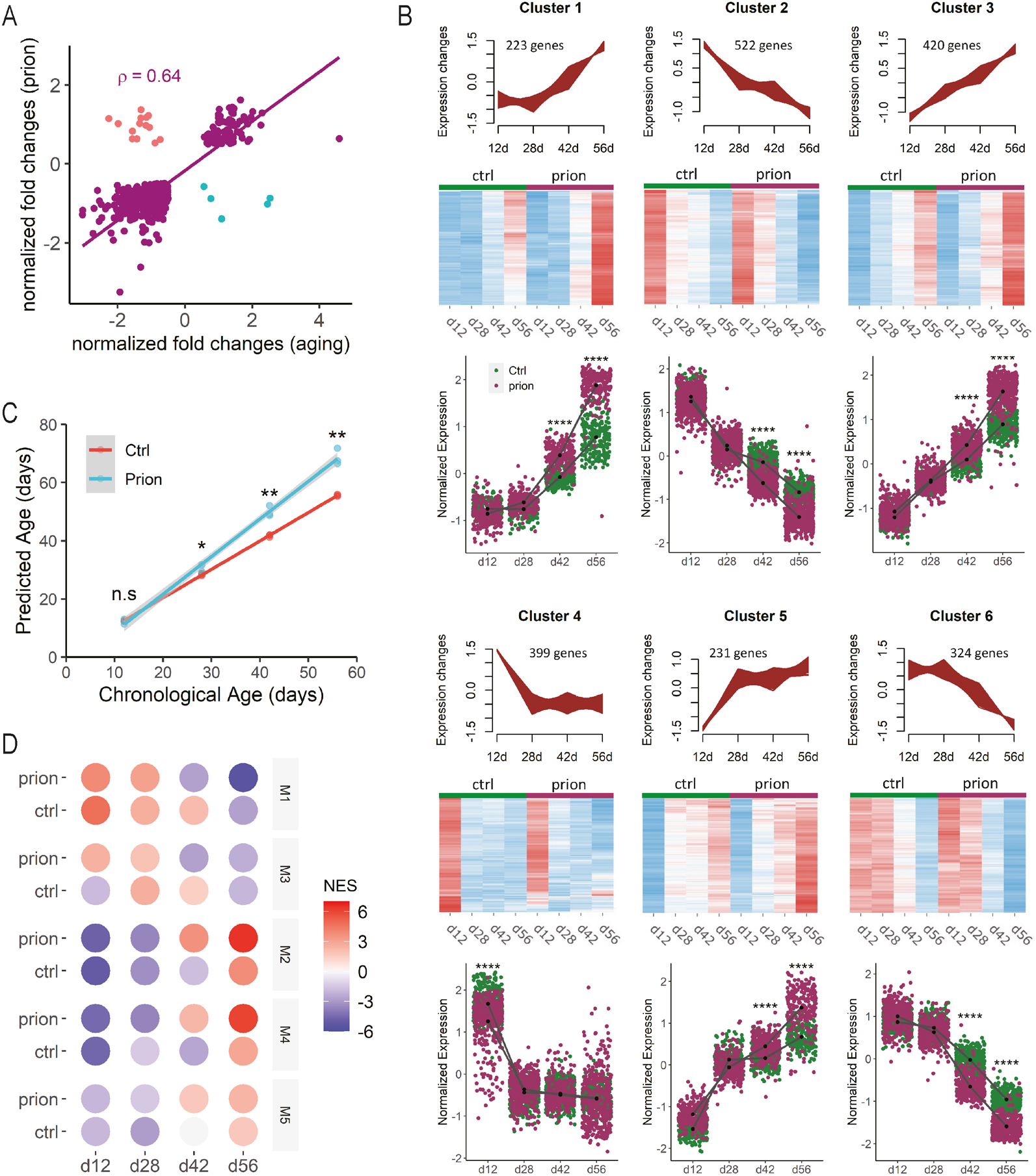
Prions accelerate biological aging in COCS. **A**, Normalized fold changes of DEG commonly identified in prion-exposed and naturally aged COCS. ρ: Spearman’s correlation coefficient. Genes show different directions of changes in the two conditions are highlighted in cyan or red. **B**, Upper panel of each cluster: graphs showing the aging-associated gene clusters classified by fuzzy c-means. Only those genes with cluster membership values > 0.5 are shown. Middle panel for each cluster: heatmaps showing the relative expression levels of clustered genes in the presence and absence of prions. Bottom panel for each cluster: quantifications of normalized expression levels of clustered genes in the presence and absence of prions. ****: p < 0.0001. **C**, Linear regression of chronological and predicted biological ages of NBH-(Ctrl) and prion-exposed COCS at different time points. Shading indicates 0.95 confidence interval. *: p < 0.05, **: p < 0.01. n.s: not significant. **D**, Heatmap showing the transcriptional activities of the five gene co-expression network-modules in NBH-(Ctrl) and prion-exposed COCS over time. NES: normalized enrichment score of gene set enrichment analysis.

To test this hypothesis, we selected 3104 aging-related genes through combining DEG identified by comparing control COCS at any two time points, and examined their expression changes in the presence and absence of prions. Using fuzzy c-means clustering (Bezdek, Ehrlich et al. 1984), we classified the aging-related genes into six clusters (**Fig. 3B**). Genes in clusters 2 and 6 showed downregulation over time (**Fig. 3B**), and were enriched for neuronal genes (**Supplementary fig. 4A**) and pathways associated with synaptic function (**Supplementary fig. 4B**). Genes in clusters 1 and 3 showed upregulation over time (**Fig. 3B**), and were enriched for Bergmann glia, astrocyte and microglia specific genes (**Supplementary fig. 4A**), and pathways related to interferon and inflammatory responses (**Supplementary fig. 4B**). Interestingly, genes in cluster 4, which were associated with cell adhesion and extracellular matrix (**Supplementary fig. 4B**) and exhibited strong early downregulation (**Fig. 3B**), were almost exclusively enriched in microglia (**Supplementary fig. 4A**), indicating possible coexistence of several distinct microglia populations. The cluster 5 genes showed strong early upregulation (**Fig. 3B**), and were enriched for genes highly expressed in oligodendrocyte linage cells (**Supplementary fig. 4A**). We examined how prion exposure influenced the transcriptional evolution of these aging-associated genes over time at the cluster level. Strikingly, we found that the temporal signatures of all the six aging-associated clusters were largely preserved after prion exposure; however, the overall temporal kinetics of both the upregulated and downregulated clusters were strongly accelerated in the presence of prions (**Fig. 3B**).

To further investigate the prion-dependent aging acceleration, we trained an age-predictive machine-learning algorithm based on ridge regression (Hoerl and Kennard 1970) using the time-course gene expression profiles in the control COCS, and calculated the biological ages of COCS exposed or not exposed to prions. The algorithm-predicted biological ages for the control COCS were similar to their chronological ages at all time points (**Fig. 3C**); however, the algorithm-predicted biological ages for the prion-exposed COCS were notably older compared to their chronological ages at three out of the four time points (**Fig. 3C**). Specifically, we found the biological ages of the prion-exposed COCS were approximately 3, 8 and 13 days older compared to the control COCS at day 28, day 42 and day 56, respectively (**Fig. 3C**). These data further support the conclusion that the biological aging process is accelerated in COCS after prion exposure.

To explore the mechanisms behind the accelerated aging in the prion-exposed COCS, we did a gene co-expression network analysis across all experimental conditions, and identified five distinct co-expression modules (**Supplementary fig. 5**). We evaluated the module activities across samples using gene-set enrichment analysis, and found all the five modules showed strong associations with both aging and prion exposure (**Fig. 3D**). Genes in modules 2, 4 and 5 showed time-dependent downregulation while genes in modules 1 and 3 showed time-dependent upregulation (**Fig. 3D**), all of which were drastically enhanced in the presence of prions (**Fig. 3D**). These results suggest that prions and aging essentially activate the same molecular programs.

### Anti-PrP antibody POM2 abolishes accelerated aging in prion-exposed COCS

To further examine the accelerating effects of prions on the aging process, we treated the prion-exposed COCS and the control COCS with the anti-PrP antibody fragment FabPOM2 (Sonati,Reimann et al. 2013), and sequenced their transcriptomes at 12 and 56 days. FabPOM2 treatment strongly reduced prion levels in the prion-exposed COCS (**Fig. 4A-B**). Strikingly, we found although FabPOM2 did not alter the expression of aging-related genes in the absence of prions (**Supplementary fig. 6A**), it completely abolished the accelerated aging signatures in the prion-exposed COCS (**Fig. 4C-D**). We then calculated the biological ages of the FabPOM2 treated COCS with our machine-learning algorithm, and found there was no difference between the control and the prion-exposed COCS anymore (**Fig. 4E**). Crucially, the biological ages of FabPOM2-treated COCS were similar to their chronological ages no matter whether they had been exposed to prions or not (**Fig. 4E**). These data indicate that acceleration of biological aging in the prion-exposed COCS was strictly dependent on prion replication.

**Figure 4.**
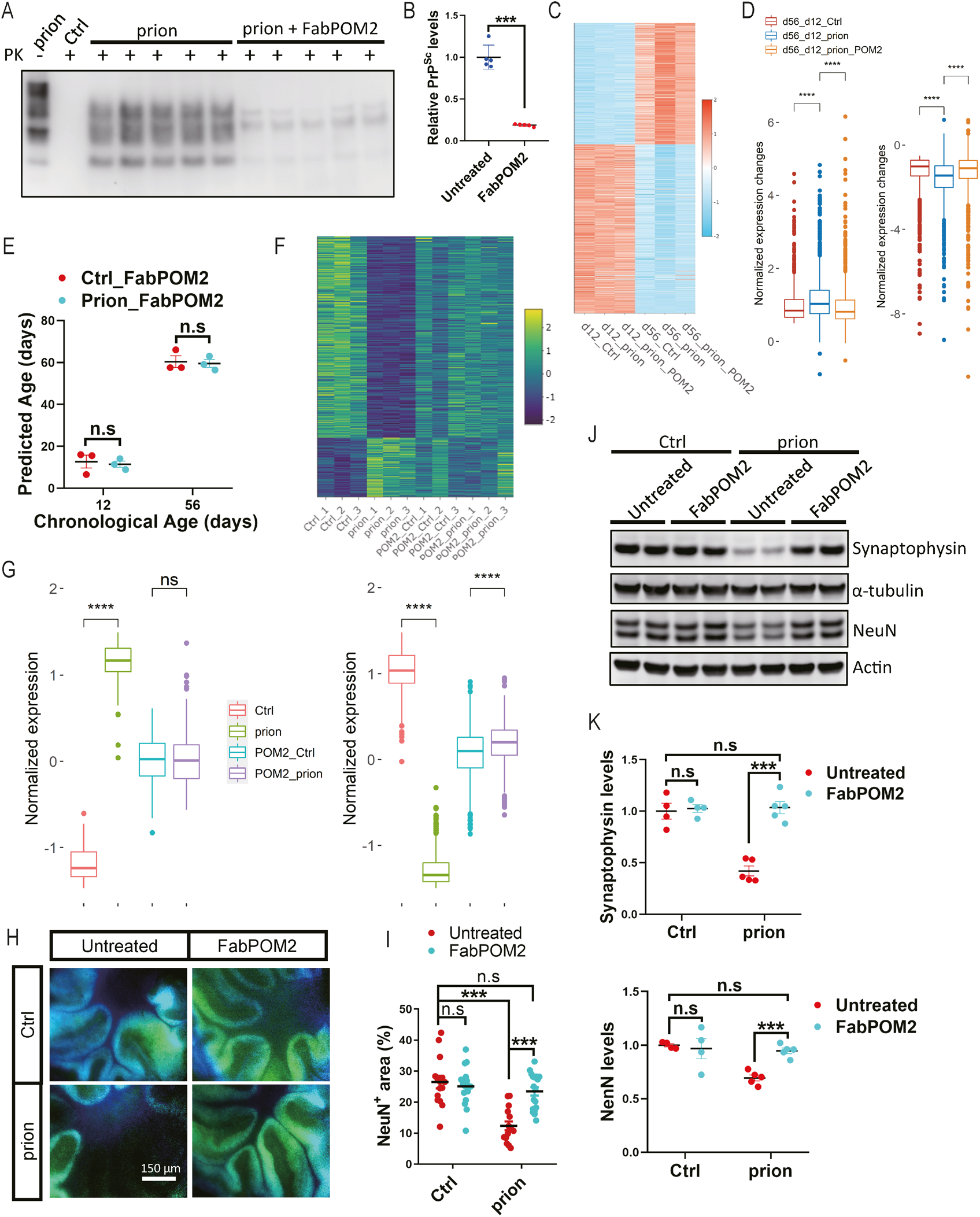
FabPOM2 treatment abolishes prion-induced aging acceleration and neurotoxicity. **A-B**, Representative western blots (**A**) and quantification (**B**) of PrP^Sc^ in prion-inoculated and control (NBH-inoculated) COCS with or without FabPOM2 treatment. ***: p < 0.001, n.s.: not significant. **C**, Heatmap showing the expression levels of aging-associated genes in NBH-(Ctrl) and prion-exposed COCS with or without FabPOM2 treatment at day 12 and day 56. Values are normalized row wise. **D**, Quantifications of normalized expression changes of the aging-associated genes shown in C between day 12 and day 56 in NBH-(Ctrl) and prion-exposed COCS with or without FabPOM2 treatment. ****: p value < 0.0001. **E**, Chronological and predicted biological ages of COCS in NBH-(Ctrl) and prion-exposed COCS treated with FabPOM2. n.s: not significant. **F**, Heatmap showing the expression levels of differentially expressed genes in prion-exposed COCS at day 56 with or without FabPOM2 treatment. Data are normalized row wise. **G**, Quantifications of normalized expression levels of genes shown in F. Left panel: upregulated genes in prion-exposed COCS. Right panel: downregulated genes in prion-exposed COCS. ****: p < 0.0001. n.s: not significant. **H**, Representative immunofluorescent images of NeuN staining in NBH-(Ctrl) and prion-exposed COCS at day 56 with or without FabPOM2 treatment. **I**, Quantifications of NeuN positive areas shown in H. ***: p < 0.001. n.s: not significant. **J**, Representative western blots showing synaptophysin and NeuN protein levels in NBH-(Ctrl) and prion-exposed COCS at day 56 with or without FabPOM2 treatment. **K**, Quantifications of synaptophysin and NeuN protein levels shown in J. ***: p < 0.001. n.s: not significant. Raw images of all western blots shown in this figure are included in supplementary fig. 7.

We then examined how FabPOM2 treatment affected prion-induced neurotoxicity. By examining the RNAseq data, we found FabPOM2 treatment completely abrogated all prion-induced molecular changes in COCS (**Fig. 4F-G**; **Supplementary fig. 6B**). Quantification of the neuronal marker NeuN through immunofluorescence and western blotting suggested that FabPOM2 treatment also abolished prion-induced neurodegeneration (**Fig. 4H-K**). In addition, we found that the progressive loss of the synaptic marker synaptophysin after prion exposure was fully prevented by FabPOM2 (**Fig. 4J-K**). These data suggest that aging acceleration by prions may be a driver of neurotoxicity.

### Accelerated biological brain aging in mouse models and patients of prion diseases

Since the progression of prion diseases is much faster than natural aging in humans and animal models, it is difficult to directly study the influence of prions on the aging process in vivo. However, if prions also accelerate brain aging in vivo, genes whose expression change in advanced age may be altered prematurely in young prion-inoculated animals or exhibit stronger changes in prion disease patients compared to age-matched control subjects.

To test this, we extracted a set of genes significantly dysregulated in the 24-month mouse hippocampus compared to 3-month (Stilling, Benito et al. 2014), and examined their expression changes in the hippocampus of 8-month compared to 3-month in the presence and absence of prions (Sorce, Nuvolone et al. 2020). In agreement with our prediction, we found many of the genes dysregulated in the 24-month hippocampus during normal aging had already shown significant changes in the 8-month hippocampus compared to 3-month in prion-exposed mice (**Fig. 5A-B**). In contrast, changes of the same genes were barely detectable in the 8-month hippocampus compared to the 3-month hippocampus in the absence of prions (**Fig. 5A-B**). To further validate these findings, we analyzed a publicly available RNAseq dataset (Forner, Kawauchi et al. 2021) examining genome-wide gene expression in the hippocampus of wild type (WT) and 5xFAD mice across different ages. Using the same approach as in **Fig. 3B**, we identified the aging-associated genes in the WT hippocampus and classified them into six clusters using fuzzy c-means clustering (**Fig. 5C**). We then examined how the expression of genes in clusters 3 and 6, two most age-predictive clusters, changed in the prion-exposed mice and AD mice compared to their respective controls. As expected, we found genes in both clusters changed much more strongly in the hippocampus of prion-exposed mice compared to the NBH control group between 8 and 3 months of age (**Fig. 5D**). In contrast, very similar expression changes were observed in the hippocampus of WT and AD mice within a similar period of time (**Fig. 5D**). Furthermore, we found a drastic upregulation of senescence-related genes, including the cellular senescence markers and members of the senescence-associated secretory phenotype, in the brains of prion-infected mice (**Fig. 5E**). These data suggest that prions also accelerate brain aging in the mouse model of prion diseases.

**Figure 5.**
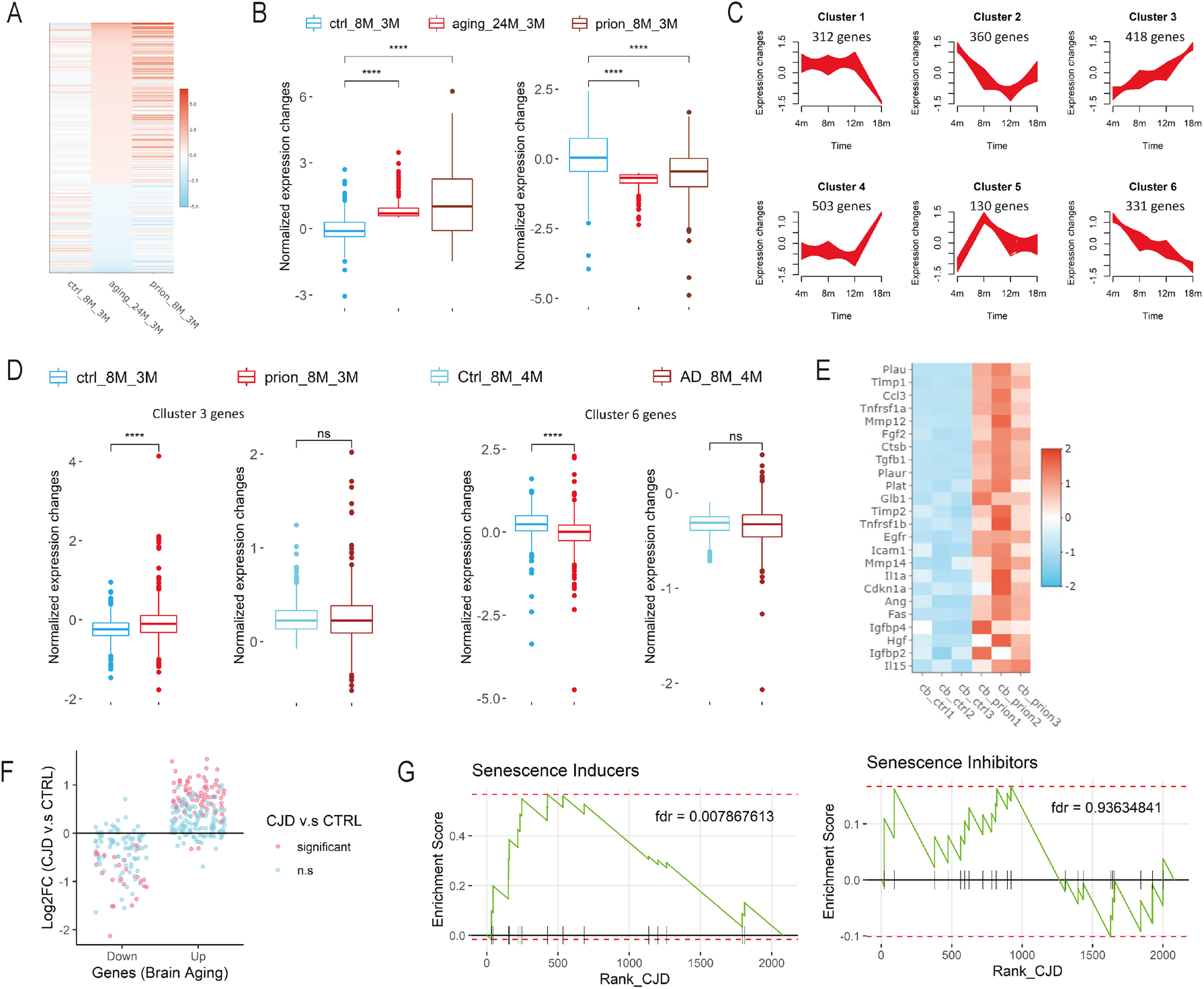
Prions accelerate biological brain aging in vivo. **A**, Heatmap showing the normalized fold changes of in vivo brain aging-signature genes across conditions. ctrl_8M_3M: 8-month vs. 3-month hippocampus in control (NBH-exposed) mice; prion_8M_3M: 8-month vs. 3-month hippocampus in prion-infected mice; aging_24M_3M: 24-month vs. 3-month hippocampus in normal mice. **B**, Quantifications of normalized expression changes of aging-signature genes shown in A. Left panel: upregulated brain aging-signature genes shown in A. Right panel: downregulated brain aging-signature genes shown A. ****: p value < 0.0001. **C**, In vivo brain aging-associated genes clustered by fuzzy c-means. Only those genes with cluster membership values > 0.5 are shown. **D**, Normalized expression changes of genes in clusters 3 and 6 shown in C in prion-inoculated mice, AD mice and their respective controls. ****: p value < 0.0001. n.s: not significant. **E**, Heatmap showing the expression levels of cellular senescence genes, including senescence markers and members of the senescence-associated secretory phenotype, in the cerebellum of NBH-(ctrl) and prion-inoculated mice. **F**, Fold changes of human brain-aging signature genes in the brains of Creutzfeldt-Jakob disease (CJD) patients compared to age-matched controls. Significant: p value < 0.05. n.s: not significant. **G**, Graphs showing results of gene set enrichment analysis of senescence inducer and inhibitor genes in the differentially expressed genes (DEG) of CJD. Statistical significance of the analysis is given as false discovery rate (fdr). DEG were ranked according to their fold changes in the CJD brains compared to age-matched controls.

To investigate whether similar biological processes are present in the brains of human prion disease patients, we examined the transcriptional changes of human brain-aging signature genes (Lu, Pan et al. 2004) in the brains of Creutzfeldt-Jakob disease (CJD) patients and age-matched control subjects (Bartoletti-Stella, Corrado et al. 2019). Strikingly, we found that most of the upregulated human brain aging-signature genes exhibited higher expression levels in the CJD brains than in the age-matched controls (**Fig. 5F**). Similarly, most of the downregulated human brain aging-signature genes exhibited lower expression levels in the CJD brains compared to the age-matched controls (**Fig. 5F**). In addition, by gene set enrichment analysis, we found the senescence-inducer genes (Avelar, Ortega et al. 2020), but not the senescence-inhibitor genes (Avelar, Ortega et al. 2020), were strongly upregulated in the CJD brains (**Fig. 5G**), indicating an induction of cellular senescence in the brains of prion disease patients. These observations suggest that possible acceleration of brain aging also present in human prion disease patients.

## Discussion

In this study, we developed a novel ex vivo model system to investigate the biological brain aging process in mammals based on COCS. This model system successfully captured the aging-modulatory effects of rapamycin, a classical anti-aging compound, validating its physiological relevance. Compared to the currently available mammalian models of brain aging, the experimental system described here is much less time-consuming and highly amenable to genetic and pharmacological perturbations. Therefore, our model system would be very useful for dissecting the complex molecular networks underlying biological brain aging in evolutionarily advanced organisms, especially when combined with high-throughput screening technologies.

Aging is a complex biological process strongly influenced by evolution. Although previous studies have identified conserved aging-related pathways across species, the pace and physiological mechanisms of aging in different species are not entirely the same (Cohen 2018). In principle, our COCS-based aging-modeling approach can be adapted to other mammalian species including primates and humans, thus providing an innovative experimental paradigm to study species-specific characteristics of biological brain aging.

Advanced age posts strong risks for developing neurodegenerative diseases associated with pathological protein aggregation (Hou, Dan et al. 2019). However, there is little consensus on why this is the case. Using our COCS-based model system, we discovered that prions strongly accelerated biological brain aging – a finding that we then corroborated in brains of experimental animals and in human CJD victims. These findings not only advance our understanding of the molecular pathology underlying prion-induced neurodegeneration, but also point to potential therapeutic interventions against prion diseases.

In essence, our data suggest that certain phenotypic manifestations of prion diseases may directly result from the accelerated changes of pathways operative in brain aging. If that is the case, it may be important to explore whether interventions that reduce certain aspects of aging might be beneficial (perhaps in combination with anti-prion therapies) in the treatment of prion diseases. Senolytic therapies may represent such candidates. Indeed, the anti-aging compound rapamycin notably suppressed prion disease pathogenesis in animal models (Cortes, Qin et al. 2012, Abdulrahman,Tahir et al. 2019). In addition, long-term injection of blood serum from young mice alleviated the clinical symptoms of prion-inoculated mice (Sorce, Nuvolone et al. 2020).

For all the novel findings discussed above, several questions surrounding are still open and need to be addressed in future studies. Firstly, previous studies have observed different effects of aging on prion disease development in animal models inoculated with prions through different routes (Avrahami and Gabizon 2011, Sorce, Nuvolone et al. 2020), and prions accumulate in the peripheral organs of animal models and human patients suffering from sporadic and variant CJD (Hill, Zeidler et al. 1997, Hilton, Fathers et al. 1998, Prinz, Montrasio et al. 2002, Glatzel, Abela et al. 2003). It would be interesting to investigate whether prions influence the biological aging process of tissues other than the brain. Secondly, since different prion strains were found to be responsible for different prion disease subtypes (Solforosi, Milani et al. 2013), future studies may determine whether the observed brain aging acceleration is a general feature of all prion strains or is restricted to the prion strains investigated in the current study.

## Materials and methods

### Animal experiment

For *in vivo* studies, adult male C57BL/6J mice purchased from Charles River Germany were used. These mice were part of the mouse cohort that had been used for our previous investigations focusing on gene expression of the hippocampus in prion diseases (Sorce, Nuvolone et al. 2020). For studies using brain slice cultures, C57BL/6J pups from the Laboratory Animal Services Center, University of Zurich were used. All animal experiments were performed according to Swiss federal guidelines and approved by the Animal Experimentation Committee of the Canton of Zurich.

### Prion inoculation

Intracerebral inoculation of adult C57BL/6J mice were performed as previously described (Sorce,Nuvolone et al. 2020). Briefly, 30 μl of 0.1% normal brain homogenate (NBH) derived from the whole brain of healthy adult CD-1 mice or same amount of prion-containing brain homogenate derived from the whole brain of terminally sick Rocky Mountain Laboratory strain of scrapie, passage 6 (RML6) infected mice was injected into the brain with 0.3-ml syringes under deep anesthesia. Health and prion disease symptoms of the inoculated mice were monitored according to a protocol approved by Animal Experimentation Committee of the Canton of Zurich.

### Organotypic brain slice culture

Brain slice cultures were prepared using cerebellar tissues from 10-12 days old mouse pups according to a previously published protocol (Falsig and Aguzzi 2008). Briefly, 350-μm thick cerebellar slices were produced using a Leica vibratome and temporarily kept in ice-cold Gey’s balanced salt solution (GBSS) supplemented with kynurenic acid (1 mM) and glucose. Slices with intact morphology were then collected and exposed to RML6, NBH or no brain homogenate for 1 hour at 4 °C. After extensive washes, six to eight slices were put on a Millicell-CM Biopore PTFE membrane insert (Millipore) and kept on slice culture medium (50% vol/vol MEM, 25% vol/vol basal medium Eagle and 25% vol/vol horse serum supplemented with 0.65% glucose (w/vol), penicillin/streptomycin and glutamax (Invitrogen)) at 37 °C in a cell culture incubator. Culture medium was changed three times per week.

### Rapamycin treatment

Rapamycin (HY-10219, MedChemExpress, USA) was dissolved in dimethyl sulfoxide (DMSO, Sigma, 472301, Switzerland), aliquot and stored at −80 °C. Freshly thawed rapamycin was added into the brain slice culture medium during medium changes from day 12 to day 56 with the final concentration of 500 nM. Same amount of DMSO was used as control.

### FabPOM2 treatment

Home-made anti-PrP antibody FabPOM2 was supplemented in the slice culture medium with 500 nM concentration from the second day after the cultures were established. The treatment continued until to the end of the experiment. Fresh FabPOM2 was added in the culture medium every time when the culture medium was changed.

### Quantitative RT-PCR

Total RNA was extracted using TRIzol (Invitrogen, USA), and reverse-transcribed into cDNA with the QuantiTect Reverse Transcription kit (QIAGEN, Germany). Quantitative RT-PCR was performed as described (Liu, Sorce et al. 2018), using the SYBR Green PCR Master Mix (Roche, Switzerland) on a ViiA7 Real-Time PCR system (Applied Biosystems, USA). The following primers were used: Mouse actin: sense, 5’-AGATCAAGATCATTGCTCCTCCT-3’, antisense, 5’-ACGCAGCTCAGTAACAGTCC-3’. Mouse Cdkn2a: sense, 5’-CGAACTCGAGGAGAGCCATC-3’, antisense, 5’-TACGTGAACGTTGCCCATCA -3’. Mouse Icam1: sense, 5’-CTGGGCTTGGAGACTCAGTG-3’, antisense, 5’-CCACACTCTCCGGAAACGAA-3’. Mouse Fgf2: sense, 5’-GGCTGCTGGCTTCTAAGTGT-3’, antisense, 5’-GTCCCGTTTTGGATCCGAGT-3’. Mouse Il1β: sense, 5’-TGCCACCTTTTGACAGTGATG-3’, antisense, 5’-AAGGTCCACGGGAAAGACAC-3’. Mouse Il6st: sense, 5’-CCAGAGCTTCGAGCCATCCG-3’, antisense, 5’-GCGCTAGCCAAATCCTTGGT-3’. Mouse Tnfrsf1a: sense, 5’-ACCAGTTCCAACGCTACCTG-3’, antisense, 5’-AGATAACCAGGGGCAACAGC-3’. Mouse Cxcl13: sense, 5’-TCTCTCCAGGCCACGGTATT-3’, antisense, 5’-TTCAGTTTTGGGGCAGCCAT-3’. Mouse Plat: sense, 5’-CAAGAGCTCAGCGTCAAGGA-3’, antisense, 5’-TTGTCTGCGTTGGCTCATCT-3’. Mouse Fas: sense, 5’-GTCCTGCCTCTGGTGCTTG-3’, antisense, 5’-AGCAAAATGGGCCTCCTTGA-3’. Mouse Mmp3: sense, 5’-CCCACATCACCTACAGGATTGT-3’, antisense, 5’-GACTGTTCCAGGCCCATCAA-3’. Mouse Mybpc3: sense, 5’-ATCCCTTTCCAGGGCAAACC-3’, antisense, 5’-CCATCATCTTGGGGTGGCTT-3’. Mouse Socs2: sense, 5’-AGCTCAGTCAAACAGGATGGT-3’, antisense, 5’-TCAATCCGCAGGTTAGTCGG-3’. Mouse Igsf3: sense, 5’-CCGGGCGAGGAATCTGC-3’, antisense, 5’-CAGCCAAAACACCCGGATTG-3’. Mouse Rims4: sense, 5’-TGGAGATTGGCTTGCAGGAG-3’, antisense, 5’-TTCCCCACACGATTACCTGC-3’. Mouse Inpp5e: sense, 5’-GCTCTAGTGCAGGGGATGTC-3’, antisense, 5’-CACCCTTGTGACGGCTTTTG-3’. Mouse Pdk4: sense, 5’-CGTACTCCACTGCTCCAACA-3’, antisense, 5’-ACACCAGTCATCAGCTTCGG-3’. Mouse Trib2: sense, 5’-AGCCCGACTGTTCTACCAGA-3’, antisense, 5’-GCTTGACACGAGTCCTCTCTT-3’. Mouse Dhx32: sense, 5’-ACCAAGAAGATGCCCGAGTG-3’, antisense, 5’-CTGTTGCAGTGTCCGTGTTG-3’. Mouse Ifi27l2a: sense, 5’-TGTTTGGCTCTGCCATAGGA-3’, antisense, 5’-TGCTGATTGGAGTGTGGCTA-3’. Mouse Vdr: sense, 5’-GCATCCAAAAGGTCATCGGC-3’, antisense, 5’-GTCGGTCTGGGGAGACAATG-3’. Mouse Scn7a: sense, 5’-TGGGGATCTTTGGAACTGGC-3’, antisense, 5’-GGCCGTGGTTCAAAGGGATA-3’. Mouse Ccl5: sense, 5’-ATATGGCTCGGACACCACTC-3’, antisense, 5’-ACTTGGCGGTTCCTTCGAG-3’. Mouse Ifitm3: sense, 5’-CTGACCATGTGGTCTGGTCC-3’, antisense, 5’-CAGGACCGGAAGTCGGAATC-3’. Mouse Cntf: sense, 5’-GCTTTCGCAGAGCAATCACC-3’, antisense, 5’-TTGCCACTGGTACACCATCC-3’. Mouse Itgb4: sense, 5’-TCTGTGCTGCAGGGAAGAGT-3’, antisense, 5’-GAGACTACCTCGGACTTGGG-3’. Mouse Cxcl5: sense, 5’-GCCCCTTCCTCAGTCATAGC-3’, antisense, 5’-AGCTTTCTTTTTGTCACTGCCC-3’. Mouse Sox11: sense, 5’-ACAGCGAGAAGATCCCGTTC-3’, antisense, 5’-CCGTCTTGGGCTTTTTGCG -3’.

### Western blotting

Western blotting was performed as previously described (Liu and Aguzzi 2020). The primary antibodies used were mouse monoclonal antibodies against actin (1:10000, Merck Millipore, MAB1501R), synaptophysin (1:5000, BD Biosciences, BD611880) and PrP (1:5000, POM1, home-made); rabbit monoclonal antibody against NeuN (1:5000, Abcam, ab177487); rabbit polyclonal antibodies against alpha-tubulin (1:5000, Proteintech, 11224-1-AP). After incubation with primary antibody overnight at 4 °C, membranes were washed three times in PBST, incubated with appropriate secondary antibody conjugated with HRP (1:10000, Jackson ImmunoResearch Laboratories) for 2 hours at room temperature. After additional washes, the membranes were developed with ECL substrates, visualized and digitized with ImageQuant (LAS-4000; Fujifilm).

### Immunofluorescence

Immunofluorescent staining of cultured brain slices was performed as described previously (Liu and Aguzzi 2020). Brain slices were fixed in 4% PFA for 30 minutes, permeabilized with 0.1% Triton X-100 in PBS and blocked with 5% goat serum in PBS overnight. After blocking, brain slices were incubated with anti-NeuN antibody (1:1000, Millipore, MAB377) for 3 days at 4 °C. After intensive washes in PBST, slices were incubated overnight with Alexa488 conjugated goat anti-mouse secondary antibody (1:3000, Jackson ImmunoResearch) at 4 °C. Stained slices were mounted on glass slides and imaged with a fluorescent microscope (Leica Biosystems).

### RNA sequencing

High throughput RNA sequencing (RNAseq) was performed as previously (Liu, Sorce et al. 2018,Sorce, Nuvolone et al. 2020). For in vivo RNAseq experiments, three mice (the entire cerebellum from each mouse as one biological repeat) per experimental condition were used at each time point. For ex vivo RNAseq experiments, cerebellar slices prepared from multiple pups from 1-3 litters with the same age were pooled together and randomly distributed into different slice culture wells with 6-8 slices per well; slices from three wells (each well as one technical repeat) per experimental condition were used at each time point. Total RNA was extracted using the TRIzol^™^ Reagent (Invitrogen, 15596-018). After cleanup with the RNeasy Plus universal mini kit (QIAGEN) and quality-check with Bioanalyzer 2100 (Agilent Technologies), sample mRNA passed the quality control was enriched with the poly-A method. Sequencing libraries were then prepared with the TruSeq RNA Sample Prep kit v2 (Illumina), and 150-bp paired-end high throughput RNAseq was performed on an Illumina Novaseq 6000 equipment at the Functional Genomic Center Zurich.

### Differential expression analysis

Differential expression analysis was done with edgeR (Robinson, McCarthy et al. 2010). Briefly, after quality control with FastQC, sequencing reads were aligned to the reference genome (Mus_musculus/GENCODE/GRCm38.p6/Annotation/Release_M23) with STAR (Dobin, Davis et al.2013). Genes with low read counts were filtered out with the default parameters in edgeR. Trimmed Mean of M-values (TMM) normalization was applied to account for compositional biases introduced by sequencing depth and effective library sizes. Significance of differential expression for each gene was tested using the QL F-test. Genes with absolute log2 (fold change) > 0.5 and false discovery rate (FDR) < 0.05 were considered differentially expressed genes (DEG). KEGG pathway enrichment analysis of DEG was performed using R package pathfindR (Ulgen, Ozisik et al. 2019).

### Clustering of aging-associated genes

Aging-associated genes were clustered according to their temporal profiles using the fuzzy c-means algorithm with the R package Mfuzz (Kumar and M 2007). Mean gene expression values (normalized counts) for each time point were normalized through logarithmic transformation and standardized with default settings. The fuzzifier parameter m and the number of clusters c were estimated with the default programs in Mfuzz. Cluster membership value > 0.5 was used for identifying the core genes in each cluster. Only the core genes in the clusters were plotted in the temporal profiles and used for downstream analyses. Pathway enrichment analysis of the clustered genes was performed using enrichR (Chen, Tan et al. 2013).

### Elastic-net and ridge regression

Age-predictive machine-learning models were trained using the R package Caret (Kuhn 2008), based on elastic-net (Zou and Hastie 2005) or ridge regression (Hoerl and Kennard 1970). Logarithmic transformed gene expression levels (normalized counts) in the control (NBH-exposed) brain slice cultures and their corresponding chronological ages were used for training the models with leave-one-out cross validation.

### Cell type enrichment of aging-associated clusters

Enrichment of core genes in each aging-associated cluster for major cerebellar cell types was performed by gene set enrichment analysis (GSEA) using the R package fgsea (Korotkevich, Sukhov et al. 2021). The core genes of each cluster were used as gene sets. The ranking list of genes for each cell type were generated based on the expression fold changes of a given gene identified in a given cell type against the rest of cerebellum (DropViz).

### Gene co-expression network analysis

Gene co-expression network analysis was performed using the R package CEMiTool (Russo,Ferreira et al. 2018). Expression levels (normalized counts) of genes detected in all experimental groups were normalized through logarithmic transformation and filtered according to variance with the default parameters. Genes in the filtered list were used for identifying gene co-expression networks with the soft-thresholding parameter beta = 20. The activity of identified co-expression modules across experimental conditions were evaluated by performing a GSEA using the genes within modules as gene sets and the median z-score values of each phenotype as rank.

### Statistical analyses

Unless otherwise mentioned, unpaired, two-tailed student’s t-test was used for comparing data from two groups, which were presented as mean ± SEM. Statistical analysis and data visualization were done using R or GraphPad Prism 8.0.

## Acknowledgement

The authors would like to thank Dr. Myvizhi Esai Selvan and Dr. Claudia Scheckel for helpful discussion, Ms. Petra Schwartz for excellent technical help in animal experiments, and Functional Genomic Center Zurich for assistance in RNAseq. A. Aguzzi is the recipient of an Advanced Grant of the European Research Council, an individual grant of the Swiss National Foundation, the Nomis Foundation and SystemsX.ch.

## Author contributions

Y. Liu designed and performed most of experiments and all bioinformatics analyses. A. Senatore and J. Guo performed RNAseq of cultured brain slices. S. Sorce and M. Nuvolone performed RNAseq of cerebellar tissues. Z. H. Gümüş provided helpful suggestions on data visualization. A. Aguzzi supervised the study and acquired funding. Y. Liu and A. Aguzzi wrote the manuscript.

## Data availability

Raw RNAseq data will be deposited on a public depository. The accession number will be available before publication. Other data have been included in the manuscript.

## Code availability

Code used for bioinformatics analyses is available from the authors upon reasonable request.

## Competing interests

The authors declare no competing interests.

---

**Supplementary figure 1.**
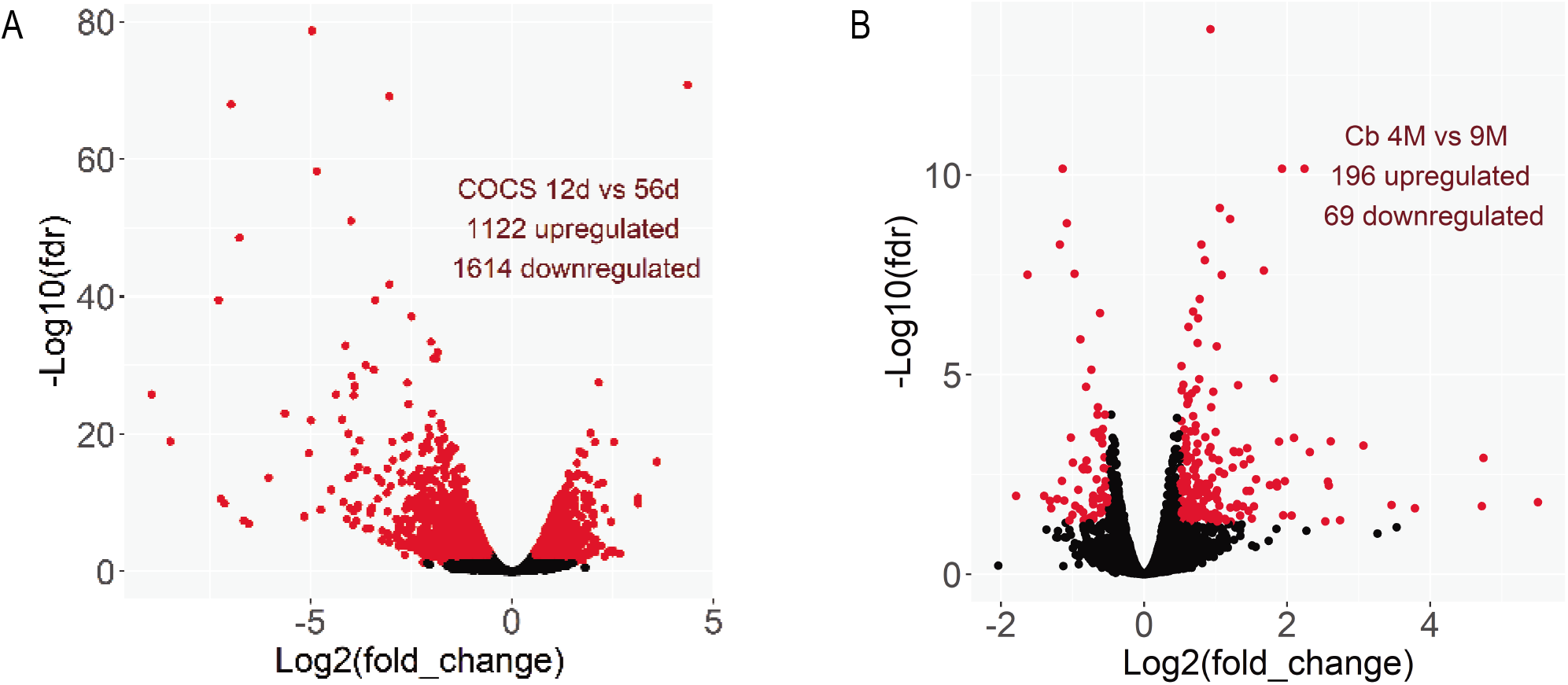
A, Volcano plot showing the dysregulated genes between the 12-day-old and 56-day-old control COCS. B, Volcano plot showing the dysregulated genes between the 4-month-old (4M) and 9-month-old (9M) NBH mouse cerebellum (Cb).

**Supplementary figure 2.**
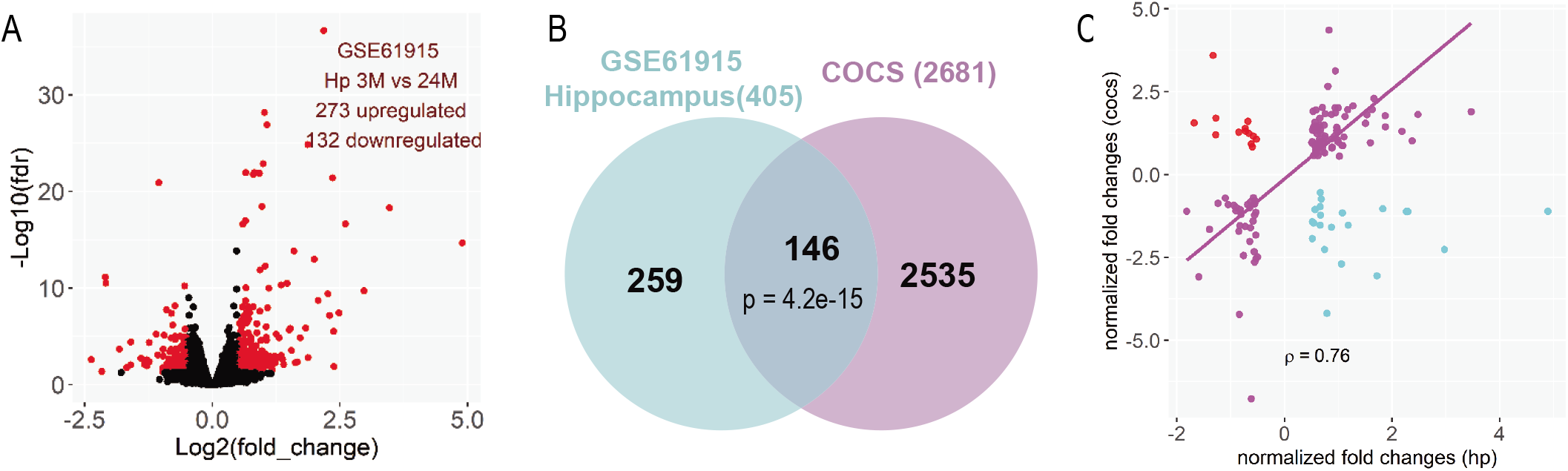
A, Volcano plot showing the dysregulated genes between the 24-month-old and 3-month-old mouse hippocampus (Hp). B, Venn diagram showing the overlaps between DEG in the aged COCS and the 24-month-old mouse Hp. p: hypergeometric test. C, Normalized fold changes of common DEG in aged COCS and the 24-month-old mouse Hp. ρ: Spearman’s correlation coefficient. Genes show different directions of changes in the two conditions are highlighted in cyan or red.

**Supplementary figure 3.**
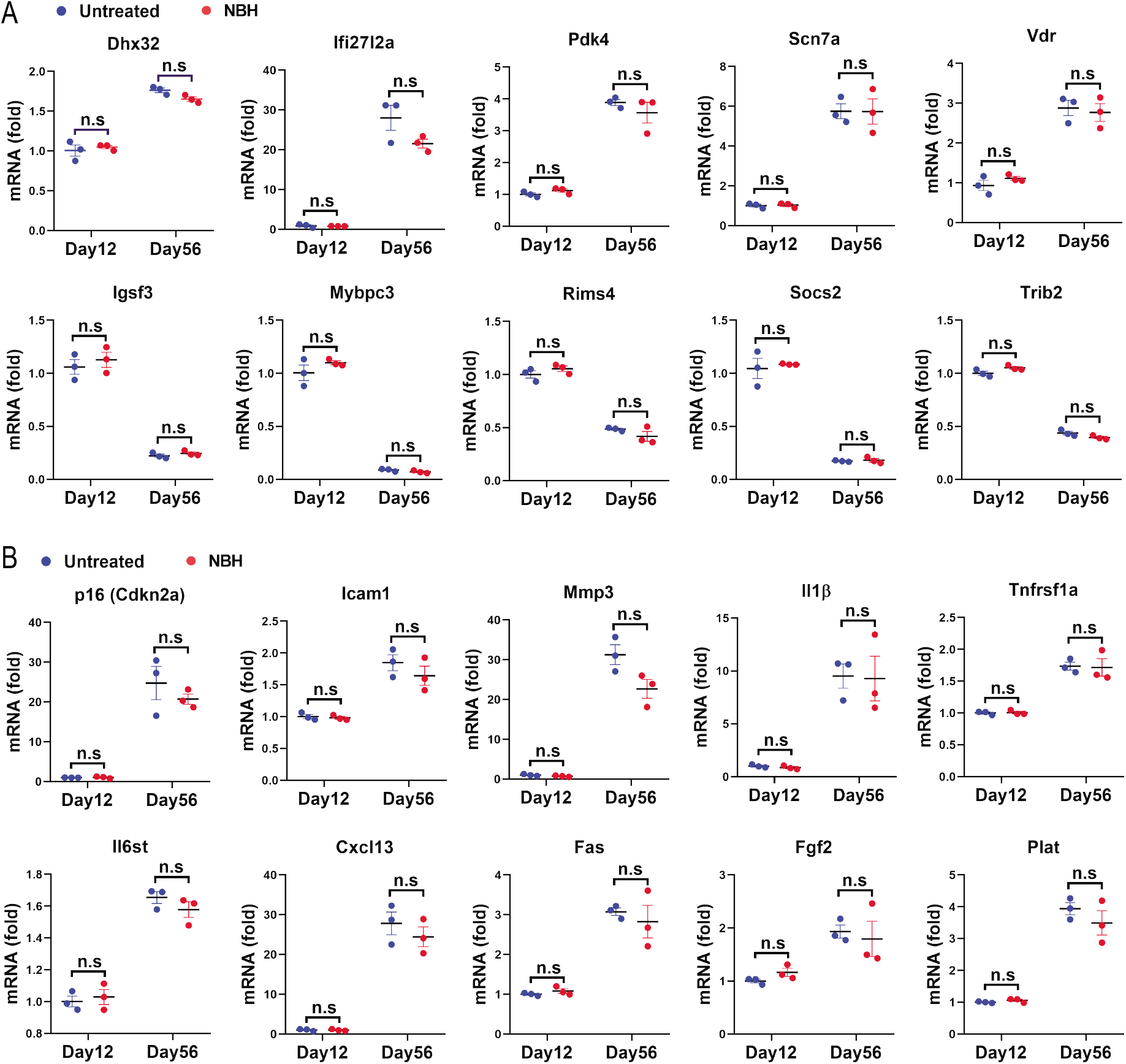
A, RT-PCR results showing similar expression levels of randomly picked age-predictive genes in naïve (untreated) and NBH-exposed COCS at day 12 and day 56. n.s: not significant. B, RT-PCR results showing similar expression levels of cellular senescence-associated genes in naïve (untreated) and NBH-exposed COCS at day 12 and day 56. n.s: not significant.

**Supplementary figure 4.**
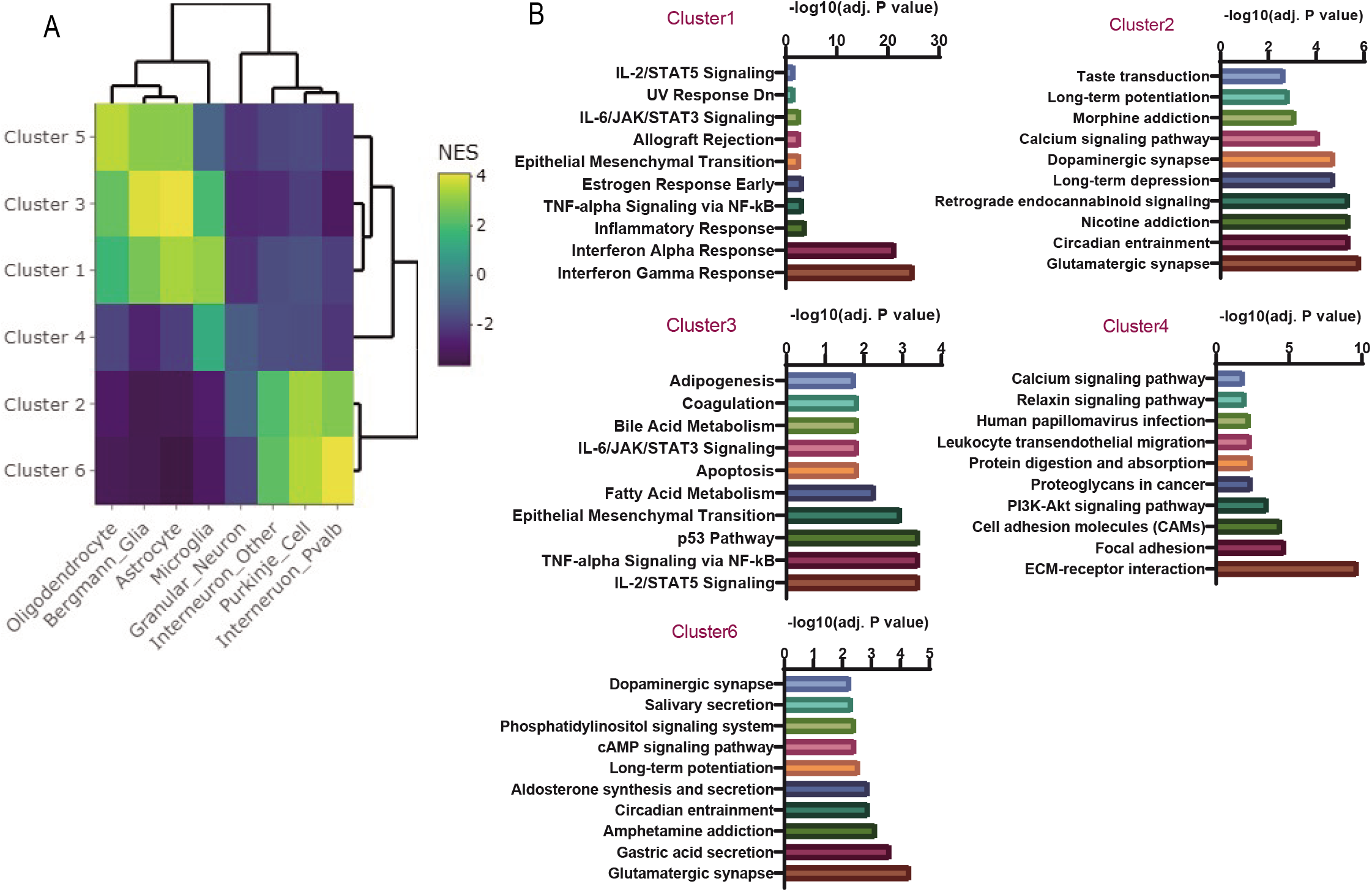
A, Heatmap showing the normalized enrichment score (NES) of gene set enrichment analysis of genes from the six clusters shown in fig. 3B for major cell types of the adult mouse cerebellum. B, Top 10 enriched pathways (adjusted p values < 0.05) for genes from the six clusters shown in fig. 3B. No pathways were significantly enriched for genes in cluster 5.

**Supplementary figure 5.**
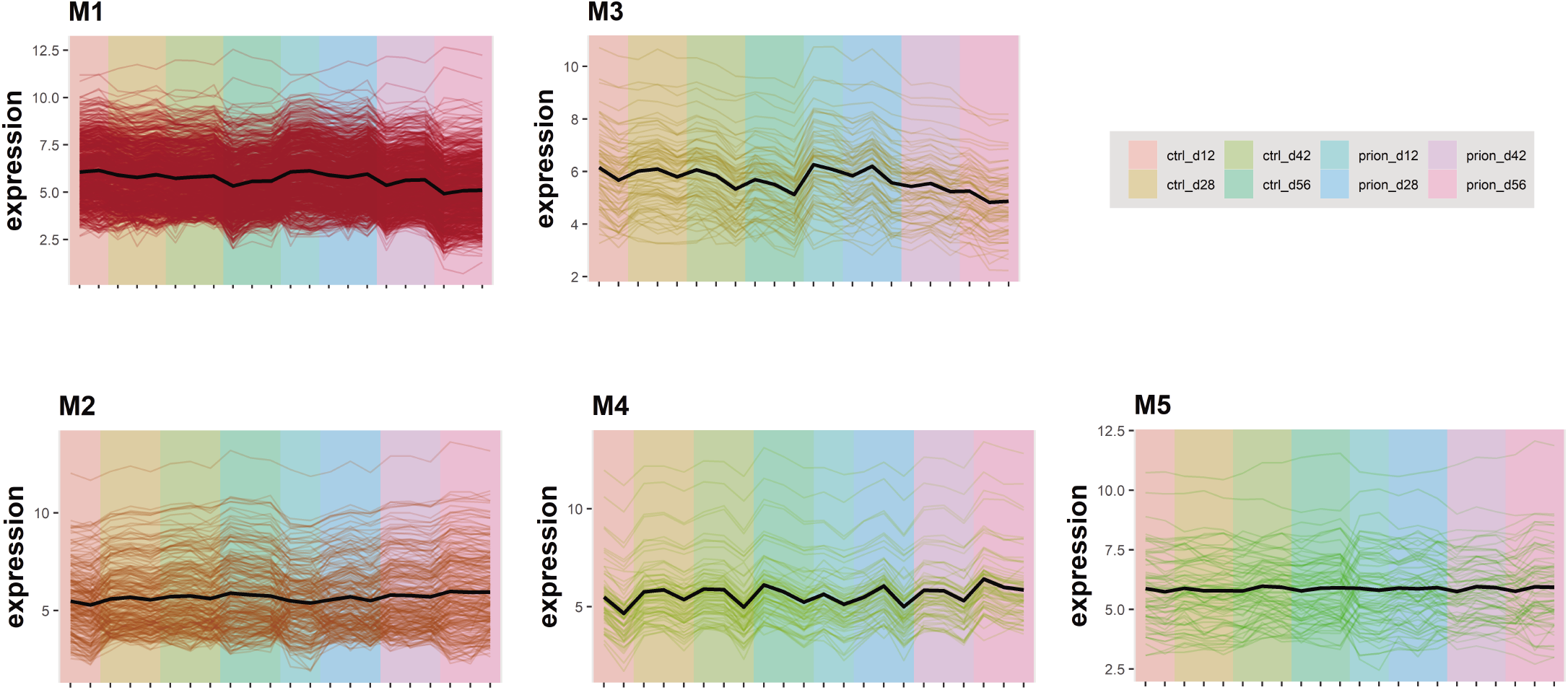
Temporal expression profiles of genes in the five gene co-expression network modules identified across experimental conditions.

**Supplementary figure 6.**
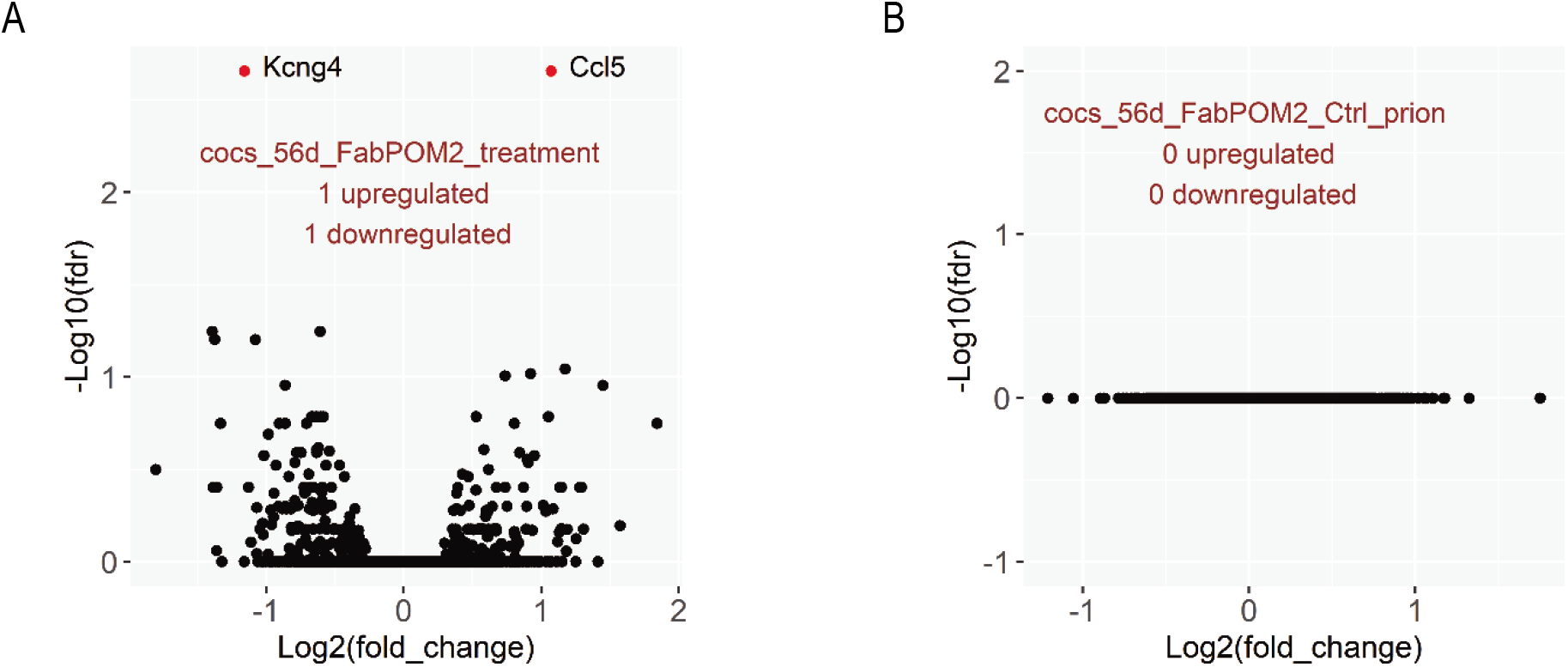
A, Volcano plot showing the only two differentially expressed genes (DEG) between FabPOM2 treated and untreated COCS at day 56 in the absence of prion infection. B, Volcano plot showing no DEG between the prion- and NBH (Ctrl) COCS treated with FabPOM2.

**Supplementary figure 7.**
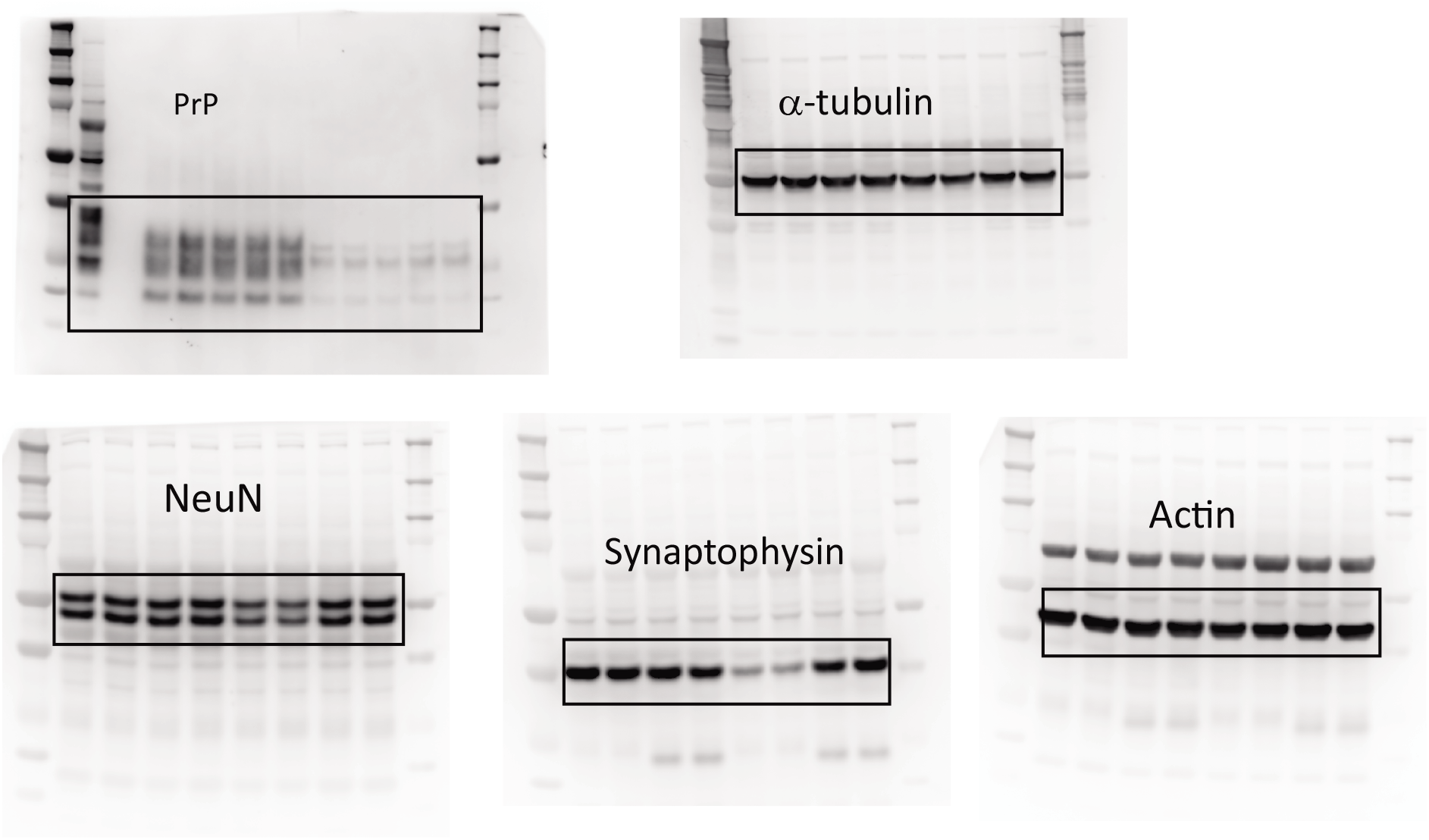
Raw images of western blots shown in Figure 4.

## Notes

### Competing Interest Statement

The authors have declared no competing interest.

